# Mispitools: An R package for comprehensive statistical methods in Kinship Inference

**DOI:** 10.1101/2024.08.16.608307

**Authors:** Franco L. Marsico

## Abstract

The search for missing persons is a complex process that involves the comparison of data from two entities: unidentified persons (UP), who may be alive or deceased, and missing persons (MP), whose whereabouts are unknown. Although existing tools support DNA-based kinship analyses for the search, they typically do not integrate or statistically evaluate diverse lines of evidence collected throughout the investigative process. Examples of alternative lines of evidence are pigmentation traits, biological sex, and age, among others. The package **Mispitools** fills this gap by providing comprehensive statistical methods adapted to a holistic investigation workflow. **Mispitools** systematically assesses the data from each investigative stage, computing the statistical weight of various types of evidence through a likelihood ratio (LR) approach. It also provides models for combining obtained LRs. Furthermore, **Mispitools** offers customized visualizations and a user-friendly interface, broadening its applicability among forensic practitioners and genealogical researchers.

## Introduction

The search for missing persons involves several steps, including preliminary investigation, archaeological research, DNA sample collection, DNA profile analysis, statistical evaluation of evidence, and communication of results (Puerto et al., 2021). The main aim of the search is to link two entities, the Missing Person (MP) and the Unidentified Person (UP). The MP represents an identity without a body. The UP refers to a body without identity, whether it corresponds to an individual whose biological identity is unknown, who is alive, or to unidentified human remains that are dead. In recent years, Forensic Investigative Genetic Genealogy (FGG) has been used to search for missing persons (Kling et al., 2021). In this area, dense SNP data are used in large databases to identify distant relatives of MPs (beyond first cousins). Historical records are combined with results from genetic analyzes to assist in the search. This approach is also implemented in criminal investigations, where UP genetic data could correspond to DNA present in a crime scene, suspected to be from the perpetrator.

During the search, genetic and non-genetic data from both entities are collected as evidence. Testimonies, historical records, judicial documentation, interviews with families, and dental and medical records are some of the evidence that contribute to the MP’s data, collected during the preliminary investigation. Generally, due to the absence of DNA data for MP, genetic information from relatives is used to perform a kinship test. Data about UP are collected from unidentified human remains and the surrounding context of the remains. For example, cemetery books could provide the date of inhumation and sometimes the date of death. In addition, different forensic techniques can contribute to estimating the time since death (Calla et al., 2021) and clues related to the crime scene, such as the analysis of bloodstain patterns and the estimation of the time since deposition (Vale et al., 2023). In some specific cases, such as child abductions and related cases, the UP is a living person who requires the proof of his biological identity (Puerto et al., 2021; Marsico et al., 2021).

In a DNA-database search, each UP is compared to the MP’s relatives using a statistical DNA kinship test based on the likelihood ratio (LR). A potential match is declared if the LR exceeds a specified threshold *T* (Kruijver et al., 2014). With recent advances in DNA genotyping and database search software, forensic professionals have incorporated DNA database search as a common practice in human identification cases. This area has been the target of multiple developments and guidelines to address opportunities and challenges. More recently, different methodologies such as Open Source Intelligence Techniques and social network analysis have allowed, in some cases, the rapid construction of preliminary investigation databases used in forensic settings (Dincelli et al., 2023). However, opportunities and challenges for these databases have not been studied as thoroughly as those of DNA databases. Moreover, despite their crucial importance in identifications, only some approaches mathematically formalize the possibility of non-genetic data, usually gathered during the preliminary investigation step, in the search process (Marsico and Caridi, 2023).

Typically, results obtained in any step of the search could lead to more investigation in another. For example, in a case where the work is done with samples from burned remains, with insufficient DNA data, DNA-database searches are hampered by low statistical power. Statistical power refers to the probability of reaching a conclusion when testing a specific hypothesis with the available data (Kling et al., 2017). More data must be collected if the statistical power is deemed too low. For example, this can be done by recruiting additional family members (Vigeland et al., 2020). This implies doing more research in the preliminary investigation step to find more relatives in the familial pedigree. In addition, low statistical power usually leads to a high number of false positives. This implies the need to rationally select a subset of UPs (potential matches) to collect more genetic data (Marsico et al., 2021). In these cases, other information collected during the preliminary investigation, such as biological sex, age, and pigmentation traits, becomes also useful. Moreover, there are cases where genetic data alone are not enough to reach a conclusion. Such cases include paternity tests where DNA-based kinship can establish a parent-offspring relationship, but not the direction; that is, it cannot resolve who is the parent and who is the child. This is a common situation in mass graves and in FGG. Another example involves individuals found to be siblings, but based on DNA data, we cannot distinguish the younger from the older. These aspects elucidate the requirement for an integrative approach when considering missing persons search, where different lines of evidence must be taken into account.

On the Comprehensive R Archive Network (CRAN), there are several packages to compute kinship testing and statistical power. We describe some of them: **Familias** comprises the open-source R code of the widely used Familias software (Kling et al., 2014). **dvir** package is oriented towards computing kinship testing in disaster victim identification cases (Vigeland, 2021). **pedprob** package allows for computing genetic marker probabilities and pedigree probabilities by implementing the Elston-Stewart algorithm (Elston and Stewart, 1971). **forrel** is a package with functions oriented toward missing person cases (Vigeland et al., 2020). **forensit** and **fbnet** offer an information-theoretic approach for kinship testing (Marsico et al., 2024; Chernomoretz et al., 2022). **kinship2** package allows for computing the kinship coefficient matrix and provides useful functions for the pedigree plot (Sinnwell et al., 2014). However, there is a lack of statistical tools for incorporating other, non-genetic, data into the search workflow.

In this paper, we introduce our R package of missing person identification tools, named **mispitools**. This package implements methods for computing the LR based on preliminary investigation characteristics, LR thresholds, error rates analysis, models for combining evidence, and others. These methods are based on computational simulations, also provided by **mispitools**, which allow researchers to study the expected outcomes of the search process based on a wide range of evidences (DNA, pigmentation traits, age, etc.). This package and its details are available on CRAN at https://CRAN.R-project.org/package=mispitools. The **mispitools** functions implement the methods described previously in several publications (Marsico et al., 2021; Marsico and Caridi, 2023; Marsico et al., 2024; Marsico and Egeland, 2024), and new functions have also been incorporated. Some of these methods have been developed to deal with specific contexts, such as DNA-based missing person identification cases hampered by low statistical power. Here we show a wider applicability of the methods, extending the field in which they can be used. Importantly, we show how to combine approaches that previously haven’t been jointly considered, elucidating their utility in examples. Last but not least, all the methods described have been used in real cases, so this package also summarizes the cumulative experience in the missing person search field.

The paper is structured as follows. First, we introduce the methodology regarding the search process for missing persons, DNA kinship tests, likelihoods calculations for non-genetic data, error rates, LR threshold selection, and performance metrics. Then, we describe the **mispitools** package and illustrate its use in two examples. Finally, we provide a summary.

## Methodology and background

In this section, the methodological background and the statistical approach are described. The main aim is to provide a gentle introduction to the key concepts. We will deal with the basic mathematical formalization required to understand the application of the methods, which we will show in the following section. For more mathematical details, we suggest that the reader consult (Marsico et al., 2021) for threshold selection; (Marsico and Caridi, 2023) and (Marsico and Egeland, 2024) for non-genetic data LR models; and (Marsico et al., 2024) for analysis of information content. All of these works present the theoretical foundations for the methods implemented in **mispitools**.

### Overview of the search

In a missing person case, data from MP and UP are collected and compared. It usually involves several steps, where different types of data are obtained. Figure 1 presents a general overview of the entire process. Generally, data collection begins with the preliminary investigation step. Preliminary investigation data can come from different sources, such as legal documentation, testimonies from relatives and witnesses, social media information, and direct observation. Some of these data, for example, pigmentation traits, can be compared between UP and MP. Others could be obtained only from one of the entities, for example, the date of disappearance for MP. For those that can be easily compared, we name their analysis PIE (preliminary investigation evidence analysis). The other cases could be useful for studying non-evident patterns that can help in the search. This package does not address these last variables; however, useful approaches can be found in (Caridi et al., 2011, 2020).

**Figure 1.**
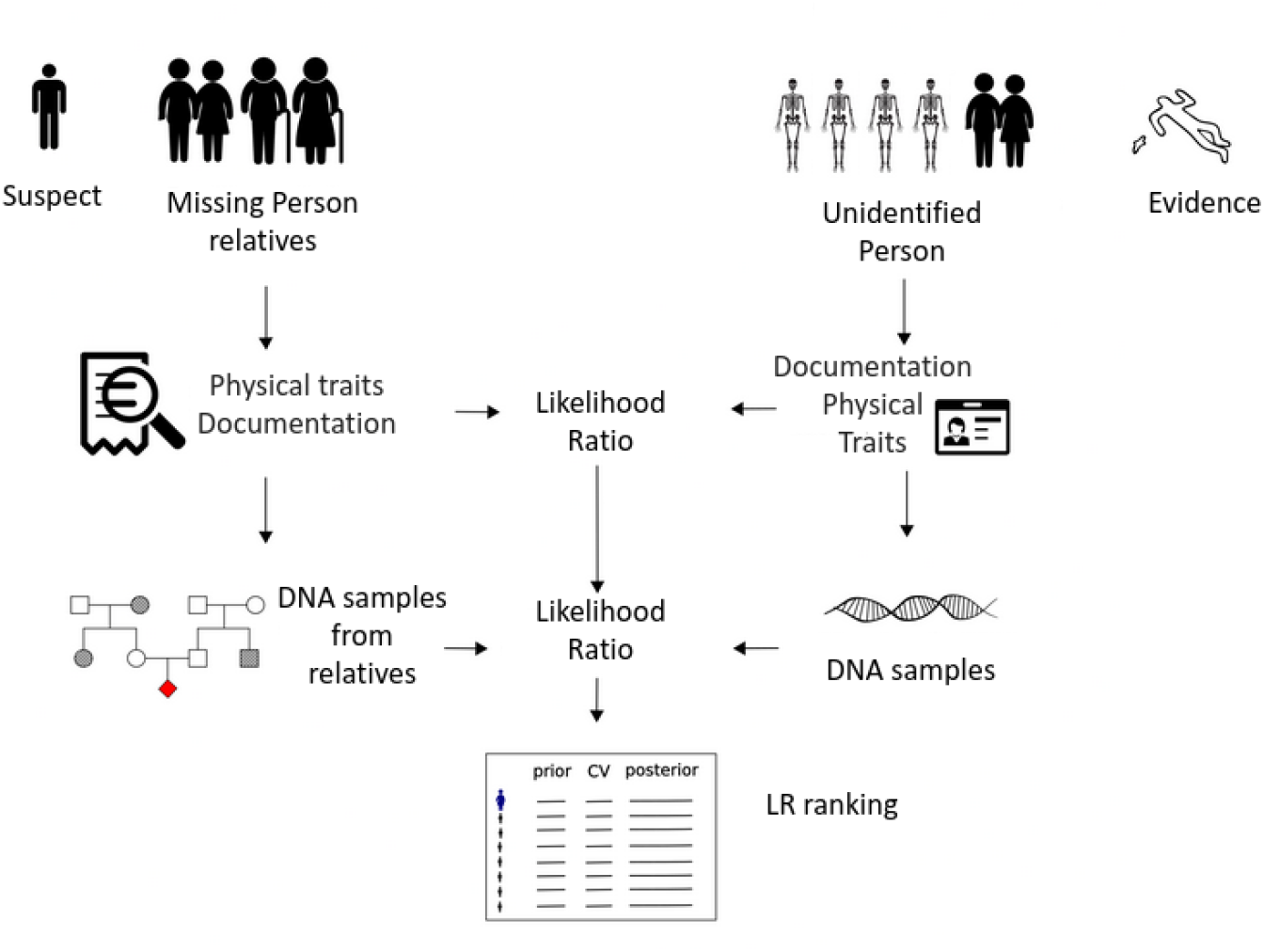
An overview of the search process for both missing person cases and criminal investigations. In missing person cases, data from unidentified persons (UP) and missing persons (MP) are collected and compared. Similarly, in criminal investigations, data from suspects (or their relatives in cases involving familial DNA searching) and evidence from the crime scene are gathered and analyzed using the same workflow. The statistical weight of both genetic evidence and preliminary investigative data (such as physical traits) is assessed through a likelihood ratio. The results are then combined to identify potential matches, generating a list of MPs or suspects who may correspond to the UP or to the individual whose traces were found at the crime scene.

**Figure 2.**
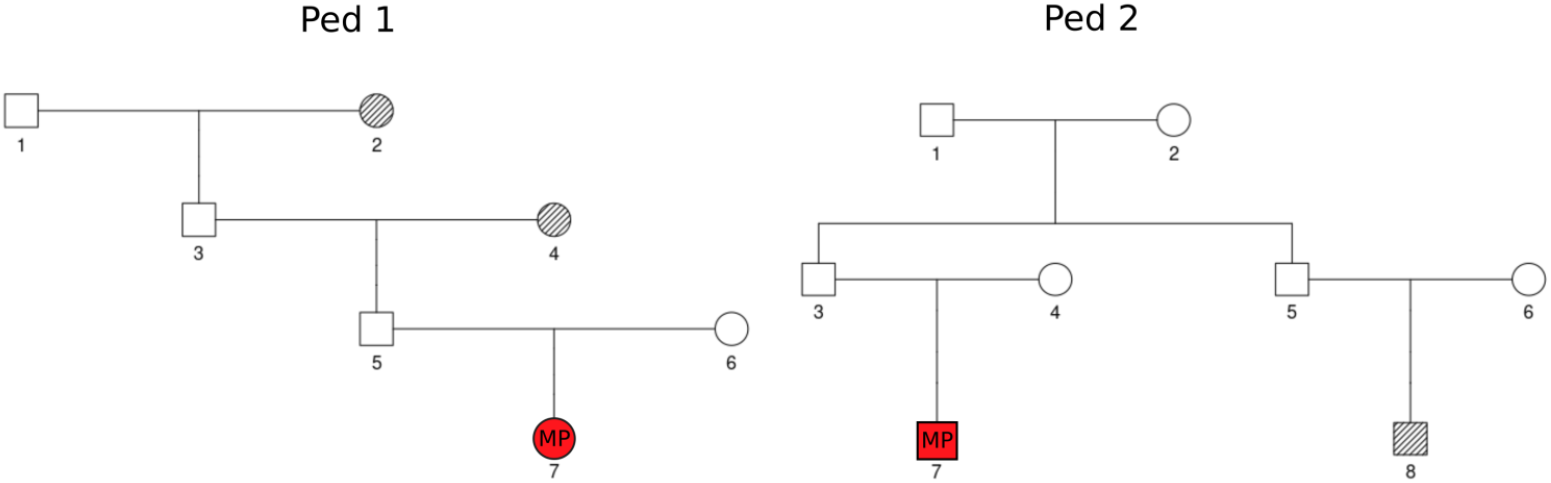
Pedigrees 1 and 2. Genotyped relatives of MP are dashed. Missing persons are in red. Id number for members is denoted.

**Figure 3.**
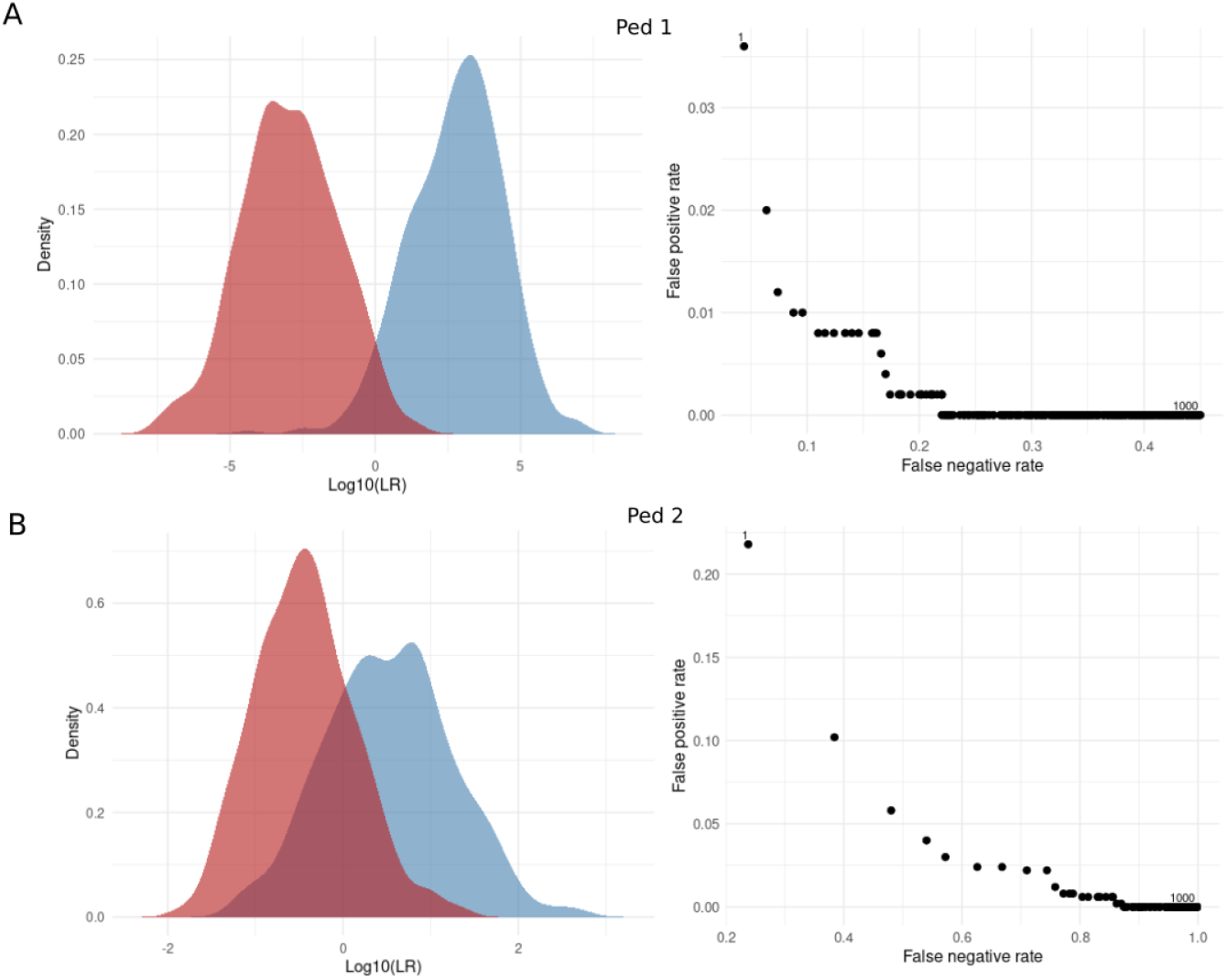
Log10(LR) distributions (left). Decision plots (right) indicate FPR and FNR for each T from 1 to 1,000. A. Present results for Ped 1, B. Corresponds to Ped 2.

During the laboratory step, through DNA analysis, genetic data from MP’s relatives (the reference pedigree) and UP are collected. DNA data consist of genetic markers for STR and/or SNP. These data can be compared using a DNA-based kinship test. We name this process FDE analysis (forensic DNA evidence analysis). In addition, in the laboratory step, techniques such as DNA phenotyping, anthropological, and entomological analysis can be used to infer traits of individuals, generally from the remains of UP (Vidaki et al., 2017). Usually, these data are compared with the preliminary investigation data from the MP.

All data collected are integrated and compared to declare an identification, both PIE and FDE. When possible, a statistical weight of the evidence is measured using the LR approach. This is the standard in genetic data, but less common in other types of evidence. Finally, results are reported.

### Statistical weight of the evidence

One important concept in forensic science is the statistical weight of the evidence. It allows us to compare the probability of observing the data given competing propositions through the LR. In missing person cases, the propositions are the following: *H*_1_ : UP is the MP, and *H*_2_ : UP is not the MP, and is a random person taken from the reference population not related to MP. The general formula for computing the LR is presented below.

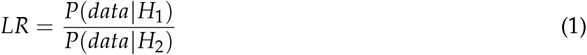

Here, *data* refers to different lines of evidence that could be mathematically formalized and compared. For example, we are searching for a missing person, named *MP*_*i*_. The color of *MP*_*i*_ ’ eyes is brown. We are analyzing a specific unidentified person (named *UP*_*j*_), whose eye color is also brown. Evaluating the statistical weight of the evidence, in this case, the color of the eye, implies computing the probability of observing the brown eye color of *UP*_*i*_, given that *UP*_*i*_ is *MP*_*i*_, and comparing it with the probability of observing the same given that *UP*_*i*_ is not *MP*_*i*_. In a simple example, we would expect that if *UP*_*i*_ is *MP*_*i*_ (*H*_1_ is true), both colors must be the same, and therefore this probability is 1. In contrast, if they are not the same person and *UP*_*i*_ is a random person from the reference population, the probability of observing the color of the eyes of *UP*_*i*_ is the frequency of brown eyes in the reference population. If that frequency is, for example, 0.5, we have

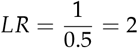

This value is interpreted as: it is 2 times more probable to observe the data given that *H*_1_ is true than if *H*_2_ is true. Intuitively, it could be guessed that a higher *LR* favors *H*_1_ against *H*_2_. In another example, both *UP* and *MP* have blue eyes. The probability of observing this, given *H*_1_, is the same. But because the blue color is less frequent, with a probability of 0.1, we have

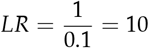

This example elucidates one of the characteristics of the search: rarer attributes can provide more statistical weight. This property has been studied through information-theoretic approaches (Marsico et al., 2024). In more realistic scenarios, the LR computation is more difficult. For PIE, different phenomena such as typing errors, uncertainty in testimonies, and technical problems in techniques such as DNA phenotyping must be incorporated into the model. For FDE, mutations, drop-in and drop-out, as well as other population genetic processes, are also incorporated. In the following subsection, specific LR models for different lines of evidence are described.

### Preliminary Investigation Evidence

We define several variables from PIE: biological sex (*S*), age (*A*), and pigmentation traits (*C*). *S* is categorized as female (*F*) or male (*M*). Age is a continuous variable from 0 to 100 years. The composite variable *C* includes hair color (*C*_*H*_), skin tone (*C*_*S*_), and eye color (*C*_*Y*_), according to the HIrisPlex system (Walsh et al., 2013). Specifically, *C*_*H*_ = 1, 2, 3, 4 (with 1 for blonde hair, 2 for brown hair, 3 for red hair, and 4 for black hair), *C*_*S*_ = 1, 2, 3, 4, 5 (representing skin tones from very pale to dark or black), and *C*_*Y*_ = 1, 2, 3 (with 1 for blue, 2 for intermediate, and 3 for brown eyes).

For *S*, we present the LR model, named *LR*_*S*_:

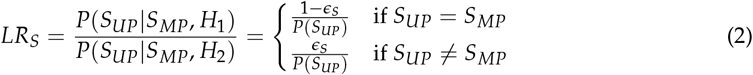

In this model, *S*_*UP*_ and *S*_*MP*_ represent the biological sexes of unidentified and missing persons, respectively. The analysis considers scenarios of sex concordance (*S*_*UP*_ = *S*_*MP*_) and discordance (*S*_*UP* ≠_ *S*_*MP*_). An error parameter, *ϵ*_*S*_, compensates for possible inaccuracies due to erroneous testimony, data entry errors, or laboratory misinterpretations.

The age variable *A* is continuous, integrating uncertainties in *A*_*MP*_ from inaccurate testimony and *A*_*UP*_ from laboratory estimates, including DNA predictions or anthropological analyzes (Vidaki et al., 2017). These uncertainties define the age ranges *A*_*UP*_ = {*UP*_min_, *UP*_max_ }and *A*_*MP*_ = {*MP*_min_, *MP*_max_}, detailed in (Cunha et al., 2009). An overlap in these ranges, indicating concordance, is measured by the Boolean variable *ψ*, where *ψ* = 1 denotes overlap and *ψ* = 0 does not overlap. The LR model for *A* is described below:

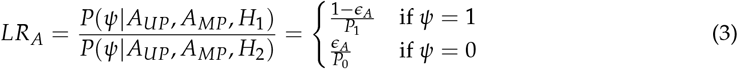

In this model, *ϵ*_*A*_ denotes the error rate of data entry and uncertainties in estimating age ranges using laboratory analyzes. *P*_1_ and *P*_0_ represent the frequencies of the *ψ* values in the reference population, where *P*_1_ is the frequency of individuals who meet the age overlap with *MP*, and *P*_0_ is the frequency of those who do not. Hence, *P*_1_ + *P*_0_ = 1.

In the case where we are dealing with a single pigmentation trait, for example, hair color (*C*_*H*_), we can avoid conditional dependency on others. Therefore, the LR is defined as follows:

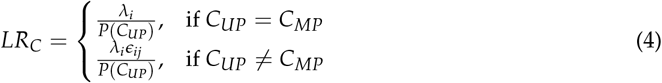

The error rate for hair color is defined for each 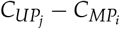 pair, reflecting the classification difficulty between colors. For instance, the error rate differs between more similar colors like blonde and red compared to distinctly different colors like black and red. A normalization constant *λ*_*i*_ ensures the total probability for a given *MP* equals 1.

For multiple pigmentation traits, the conditional dependency must be taken into account. Given the phenotypic characteristics (*C*) of an MP and UP, including hair, skin, and eye color, and the defined error rates *e*_*h*_, *e*_*s*_, and *e*_*y*_, the LR equation integrates error rates and concordance indicators (*δ*_*h*_, *δ*_*s*_, *δ*_*y*_):

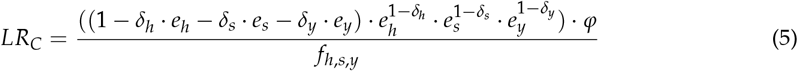

The normalization factor *φ* ensures that the probabilities sum to one and *f*_*h,s,y*_ adjusts for the observed frequency using Laplace smoothing.

### Forensic DNA evidence

For genetic evidence in Missing Persons Identification (MPI) scenarios, the genetic profile *G*_*α*_ consists of a set of markers *M*_*α,i*_, which capture the genotypes of UP and any potential relatives of MP. Using parameters such as mutation rates and population structure, the LR for this scenario is defined as follows.

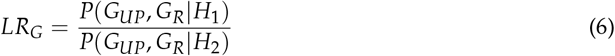

Here, {*G*_*UP*_, *G*_*R*_} represents the genotype data for UP and MP relatives, with *P*(*G*_*UP*_, *G*_*R*_ | *H*_*i*_) indicating the likelihood of observing the specific genetic profiles under hypothesis *H*_*i*_. LR quantifies the comparative likelihood of the evidence under hypothesis *H*_1_ versus *H*_2_.

### Combining evidence

We assume that genetic and non-genetic evidence are independent. Thus, for a specific scenario *a*, the combined Likelihood Ratio (LR) is modeled as:

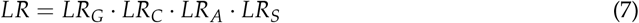

This formulation presumes the conditional independence of the genetic, color, age, and sex variables for the calculation of LR. However, it accounts for the interdependence among color traits, specifically between hair, skin, and eye color, thus refining the model to better reflect the nuances of phenotypic characteristics.

### Computational simulations

The methodology to perform simulations of both FDE and PIE is described in this section. Several approaches have previously been proposed for FDE simulations (Vigeland et al., 2020; Marsico et al., 2021; Kling et al., 2017). However, there is a lack of implementations with regard to PIE simulations. Taking into account both types of evidence, complete missing person search scenarios can be explored and different downstream analyzes can be performed using simulated data (Vigeland et al., 2020). We describe a general approach for performing simulations for PIE and FDE below, accompanied by a pseudocode for implementing these simulations.

#### Algorithm 1 Evidence Simulations

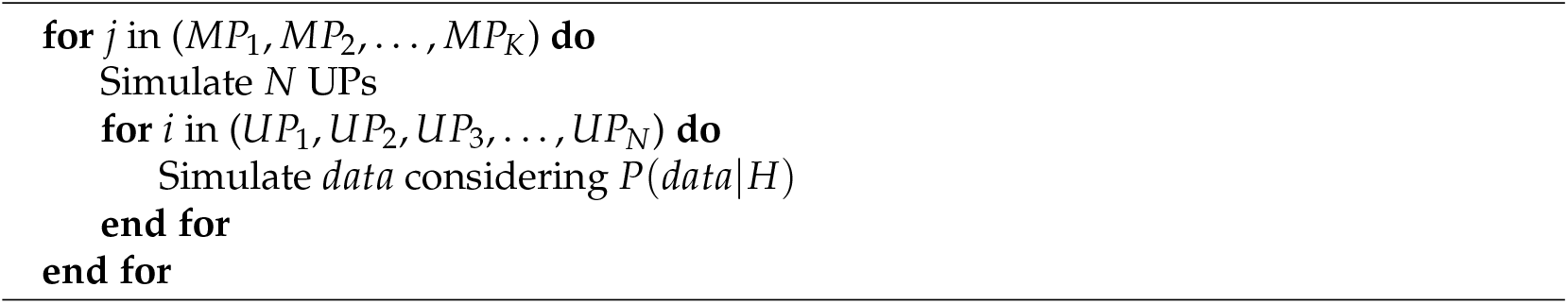

The algorithm outlines that for each *MP*_*i*_, a set of *N* UPs is simulated, and for each *UP*_*j*_, data (either genetic or nongenetic) are generated assuming hypotheses *H*_1_ or *H*_2_. Typically, simulations include 10,000 UPs for each hypothesis. As a practical example, we can simulate the sex variable *S* for an MP identified as female (*S*_*MP*_ = *F*) with an error rate *ϵ*_*S*_ = 0.05, resulting in two lists of UPs: one under *H*_1_ with an expected female proportion of 0.95 (9,500 out of 10,000), and another under *H*_2_ reflecting the frequency of the general population *P*(*F*). This method can be extended to other data types. Once simulated, the PIE and FDE data can be used to compute LRs.

### Performance metrics

In this section, we introduce performance metrics used for both FDE and PIE (and the combination of both). Performance studies allow forensic practitioners to evaluate the expected results of the search considering available data and LR models. This means that researchers can study, for example, the number of expected cases above a LR threshold. This could be used to decide between advancing the search with available data or putting more effort into gathering more evidence (Marsico et al., 2021), among other applications. Here, we define two general approaches for performance analysis, one based on computational simulations and the other on information theory metrics.

The first method utilizes the output of the simulations to obtain the *Log*_10_(*LR*) distributions considering that *H*_1_ or *H*_2_ is true. This allows defining the false negative rate (FNR) as a function of the threshold *T*,

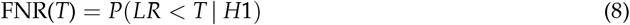

and similarly for the false positive rate (FPR):

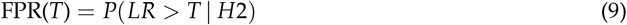

For each pedigree, with conditional simulations, we estimated FNR and FPR for each integer *T* between 1 and 1,000. The Decision Threshold (DT) approach can be used to select that subset of UPs for which gathering more FDE and PIE data could help in solving the case. DT is the value of *T* that minimizes the following weighted Euclidean distance (WED) to the theoretical optimum:

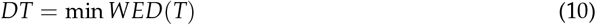

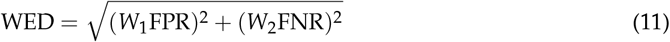

The choice of weights *W*_1_ and *W*_2_ reflects the relative importance of false positives and negatives. In this case, we used *W*_1_ = 10, and *W*_2_ = 1 (see Marsico et al. (Marsico et al., 2021) for more details). The Matthews correlation coefficient (MCC) can be employed as a summary measure considering all defined error rates.

Other metrics based on information theory (IT) have been proposed (Ramos et al., 2020). One of the main advances is that they use the conditional probability tables *P*(*data*|*H*_1_) and *P*(*data*|*H*_2_) as input, therefore, they can be directly computed avoiding computational simulations. In particular, the Kullback-Leibler metric has recently been proposed as a way to study the improvement of the addition of a new relative to a pedigree (Marsico et al., 2024). It quantifies the difference between two probability distributions, *P*_1_(*G*) and *P*_2_(*G*), as follows:

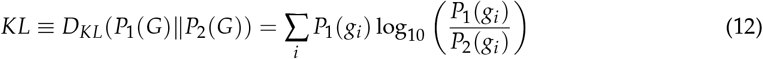

*KL* measures the expected logarithmic difference between the distributions, highlighting deviations that occur frequently in *P*_1_. It is zero when the distributions are identical, indicating that there is no evidence of impact. The asymmetry of *D*_*KL*_ suggests analyzing the reverse divergence:

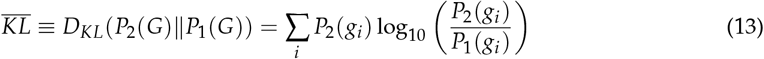

This simplifies to a sum of individual marker contributions under the assumption of independence. Metrics are expressed in dits (decimal digits), directly applying the base 10 logarithm.

### Using Mispitools

In this section, we first summarize the main functions and data structures present in **Mispitools** R Package. Then, two examples of missing person search are analyzed. The purpose of these examples is to show how **Mispitools** can be used to assist decision making in complex large-scale kinship testing cases.

### Overview of R package Mispitools

Here are some functions and their descriptions at a glance. First, functions for simulating likelihood ratios are presented.

- *simLRgen*: Calculates likelihoods ratios (LRs) based on genetic simulations. It is a function for obtaining expected LR values considering *H*_1_ or *H*_2_ as true.
- *LRsex*: Calculates LRs based on biological sex simulations. It allows for obtaining expected LR values considering *H*_1_ or *H*_2_ as true.
- *LRcol*: Calculates LRs based on pigmentation color (one-variable model) simulations. It allows for obtaining expected LR values under *H*_1_ and *H*_2_.
- *LRage*: Calculates LRs based on age simulations. It allows for obtaining expected LR values under *H*_1_ and *H*_2_.
- *combLR*: Combines LRs from different variables simulated under the indicated scenarios.
- *compute_LR_colors*: Computes LR distribution for all combinations of multiple pigmentation variables.
- *LRcols*: Calculates LRs based on multiple pigmentation color simulations. It allows for obtaining expected LR values under *H*_1_ and *H*_2_.

Other functions allow for simulating missing person search databases with both PIE and FDE.

- *makePOIgen*: Generates a genetic database for UPs. It samples genotypes from the allele frequency database.
- *makePOIprelim*: Produces preliminary investigation data for UPs. Different database models are available for different missing person search scenarios.
- *makeMPprelim*: Produces preliminary investigation data for MPs. Different database models are available for different missing person search scenarios.

Some functions allow for performance computation and LR threshold selection and performance metrics.

- *Trates*: Calculate error rates and accuracy for specific thresholds. It uses as input the simulated likelihood ratio distributions.
- *DeT*: Calculate the optimal threshold using the DT approach. It requires weights as input.
- *bidirectionalKL*: Kullback-Leibler Divergence Calculation for Genetic Markers. Calculate KL using two allele frequency databases.
- *klPIE*: Computes the Kullback-Leibler Divergence between *H*_1_ and *H*_2_ PIE likelihood matrices.

The functions presented below aim to provide graphical utilities.

- *LRdist*: Calculate the likelihood ratio based on PIE. Different models are available for different variables.
- *CondPlot*: Generates a general plot for conditioned probabilities and LR combining nongenetic variables.
- *deplot*: Displays the decision plot showing false negative and false positive rates for each LR threshold. It is an interactive chart.
- *mispiApp*: Launches a Shiny app that implements some of the core functionalities of **Mispitools**, providing a user-friendly interface.

In addition, a set of DNA databases from various countries around the world is integrated. A specialized function is implemented to manage database formats.

- *Allele frequency databases*: **Mispitools** provides a set of short tandem repeat (STR) allele frequency databases from different countries around the world. Some of these are: Argentina, China, USA, Bosnia and Herzegovina, and others.
- *getFreqs*: Allows for the use of allele frequency databases for genetic simulation. It adapts the database format.

#### Example 1. How to establish an LR threshold for each pedigree in missing person cases?

This first example aims to show how it is possible to establish an LR threshold in the DNA database search when it is hampered by low statistical power. Consider the balance between false positive and false negative rates, obtained from computational simulations. Firstly, the **mispitools** package must be installed:

~~~
install.packages(”mispitools”)
library(mispitools)
library(pedtools)
library(forrel)
~~~

Then, STRs allele frequency databases are incorporated. Also, the pedigree information is read. For pedigree construction, **pedtools** package is used. It can read .fam and .ped formats with genetic information, and also define genotypes to the members from command line. In this example, we assign genotypes to specific members in order to have reproducibility. The command for this task is presented below:

~~~
set.seed(1234)
f <-getfreqs(Argentina)
ped1 <-linearPed(3)
ped1 <-setMarkers(ped1,locusAttributes = f)
ped1 <-profileSim(ped1,N = 1,ids = c(2,4))
ped2 <-cousinPed(1)
ped2 <-setMarkers(ped2,locusAttributes = f)
ped2 <-profileSim(ped2,N = 1,ids = 8)
datasimx = simLRgen(ped1,missing = 7,numsims = 500,seed = 1234)
datasimy = simLRgen(ped2,missing = 7,numsims = 500,seed = 1234)
~~~

In the first line, the STRs allele frequency database from Argentina is loaded, with 23 STR markers. Then a two-generation pedigree structure is generated using the profileSim function. Genotypes of members of the pedigree are generated. In one case, for pedigree 1, IDs 2 (great grandmother) and 4 (grandmother) are the genotyped members. In the other case, for pedigree 2, the genotype for 8 (cousin) is defined with profileSim. Pedigrees can be plotted as follows:

~~~
plot(ped1,hatched = typedMembers(ped1))
plot(ped2,hatched = typedMembers(ped2))
~~~

`SimLRgen` allows obtaining LR values considering available genetic data in the pedigree. In this particular case, 500 genotypes for the UP are simulated considering that UP is the MP, with *H*_1_ as true. This means that genotypes compatible with available genotyped members in each pedigree are simulated. Additionally, 500 genotypes for the UP considering *H*_2_ as true are simulated. This is obtained from random sampling using the allele frequency database. For each UP genotype, the LR is computed. Therefore, 500 LR values considering *H*_1_ and 500 more considering *H*_2_ as true are obtained. These data are stored in a data frame. In the next steps, this information will be used for performance metrics.

~~~
LRdist(datasimx)
deplot(datasimx)
LRdist(datasimy)
deplot(datasimy)
~~~

LRdist plot the Log_10_ distributions when H1 or H2 is true. Deplot allows looking for false positive and false negative rates for each LR threshold from 1 to 10.000.

At this point, there are some clues for evaluating the statistical power of the pedigrees and rationally choosing an LR threshold based on error rates. However, some metrics could be computed to take a more precise decision:

~~~
DeT(datasimx, weight = 10)
“Decision threshold is: 4”
Trates(datasimx, threshold = 4)
“FNR = 0.088 ; FPR = 0.01 ; MCC = 0.904756468160616”
DeT(datasimy, weight = 10)
“Decision threshold is: 5”
Trates(datasimy, threshold = 5)
“FNR = 0.572 ; FPR = 0.03 ; MCC = 0.473596131451164”
~~~

DT computes the decision threshold based on the approach described previously (Eq. 10). It takes into account the balance between FPR and FNR. Trates specifies some metrics for the indicated threshold, such as error rates and Matthews correlation coefficient (MCC). The results show that Pedigree 1, as expected, performs better due to having closer and more relatives to the MP. However, *FPR* = 0.01 can be unmanageable in large databases. For example, analyzing 30,000 individuals could lead to 300 potential matches. Here, other information must be collected to prioritize the cases in which further genetic analyses can be performed to arrive at a conclusion.

This simple example shows how **mispitools** can be used in missing person search cases not only to compute the expected LR values considering *H*_1_ and *H*_2_, but also to provide a method to select an LR threshold. In this case, only genetic data are considered; In the following, we introduce one of the core functionalities in **mispitools**, which is its ability to compute and combine LRs from different lines of evidence.

#### Example 2. Combining genetic with preliminary investigation-based LRs

In this example, we will introduce the LR calculation for PIE and show how, combined with DNA-based LRs, search performance metrics can be improved. The first case considers only conditionally independent PIE variables. First, we generate the reference population data with the following command:

~~~
POPl <-CPT_POP(propS = c(0.5, 0.5),
               MPa = 40,
               MPr = 6,
               propC = c(0.3, 0.2, 0.25, 0.15, 0.1))
~~~

The parameter `propS` is used to indicate the proportion of biological sex, female or male. Here, we select a uniform distribution for our reference population. MPa and MPr are the age of the MP and the error range, respectively, so the age of the MP is defined between [MPa - MPr ; MPa + MPr]. Note that in this case, MP values are important for defining the population reference distribution because they are used to compute the proportion of individuals that fall within the MP’s age range and those that are outside it. The age distribution model for the reference population is uniform in this case, for simplicity, but it can be manually customized. Finally, propC indicates the proportion of one pigmentation trait (single model, without considering the dependency between multiple traits). The output of this function, stored in `POPl`, is the following:

~~~
POPl
       [,1]  [,2]    [,3]    [,4]   [,5]
F-T1 0.0225 0.015 0.01875 0.01125 0.0075
F-T0 0.1275 0.085 0.10625 0.06375 0.0425
M-T1 0.0225 0.015 0.01875 0.01125 0.0075
M-T0 0.1275 0.085 0.10625 0.06375 0.0425
~~~

The matrix obtained represents the probabilities of the phenotypes in the reference population and therefore the probability considering *H*_2_, *P*(*D*| *H*_2_), for each specific combination of characteristics. F-T1 represents a woman whose age is matched with the age of MP. F-T0 represents a woman with a mismatch in age with MP. M-T1 and M-T0 correspond to the same age categories in association with male. The numbers (columns) represent different hair colors.

To obtain the conditioned probabilities of the phenotypes considering *H*_1_ as true (*P*(*D*|*H*_1_)), we can use the following command:

~~~
MPl <-CPT_MP(MPs = “F”, MPc = 1,
             eps = 0.05, epa = 0.05,
             epc = Cmodel())
~~~

The parameters MPs and MPc indicate the biological sex and the pigmentation trait color (in this case hair), for MP. On the other hand, eps, epa and epc are the error rates. For epc, a function named Cmodel is used to define specific error rates for each pair of colors. In this case, default values are used, but they can be customized as indicated in the documentation. The output, stored in MPl, is the following:

~~~
MPl
               1            2            3            4            5
F-T1 0.877918288 8.779183e-03 4.389591e-03 8.779183e-03 2.633755e-03
F-T0 0.046206226 4.620623e-04 2.310311e-04 4.620623e-04 1.386187e-04
M-T1 0.046206226 4.620623e-04 2.310311e-04 4.620623e-04 1.386187e-04
M-T0 0.002431907 2.431907e-05 1.215953e-05 2.431907e-05 7.295720e-06
~~~

At this point, for each combination of preliminary investigation variables, we have the likelihoods of *H*_1_ and *H*_2_. Therefore, LR can be easily computed as following:

~~~
MPl/POPl
~~~

Obtaining the following output:

~~~
               1            2            3            4           5
F-T1 39.01859058 0.5852788586 0.2341115435 0.7803718115 0.351167315
F-T0  0.36240177 0.0054360266 0.0021744106 0.0072480354 0.003261616
M-T1  2.05361003 0.0308041505 0.0123216602 0.0410722006 0.018482490
M-T0  0.01907378 0.0002861067 0.0001144427 0.0003814755 0.000171664
~~~

This indicates the LR values for each combination of characteristics, considering the MP and reference population data. With these tables, each UP can be easily evaluated. For example, if we have an UP with the same characteristics as a MP (hair color = 1, female, and within the age range), we would obtain LR = 39. If we have a UP with the same hair color and age range but a different sex, the LR would be around 2. If we have a complete mismatch in the characteristics, for example, hair color = 5, outside the age range, and female, the LR would drastically drop to 0.00017.

The package **mispitools** also provides a customized plotting function to visualize likelihoods and LRs.

~~~
CondPlot(MPl,POPl)
~~~

The following plot is obtained:

These tables are useful for several applications. On the one hand, they allow us to know the expected values when we face a case where UP is MP. Also, if we consider Figure 4A, we can obtain the probabilities of each phenotype when *H*_2_ is true. If we combine these probabilities with the LR values for each characteristic (Figure 4C), we can obtain the probability distribution of LR given *H*_2_, *P*(*LR*|*H*_2_). In the same way, if we use Figure 4B, we can obtain *P*(*LR* |*H*_1_). This approach can be used to perform computational simulations and also to measure the statistical power. In fact, to directly measure the content of the information provided by the MP and reference population data, we can compute, through the klPIE function, the Kullback-Leibler divergence between *P*(*D*|*H*_2_) and *P*(*D*|*H*_1_).

**Figure 4.**
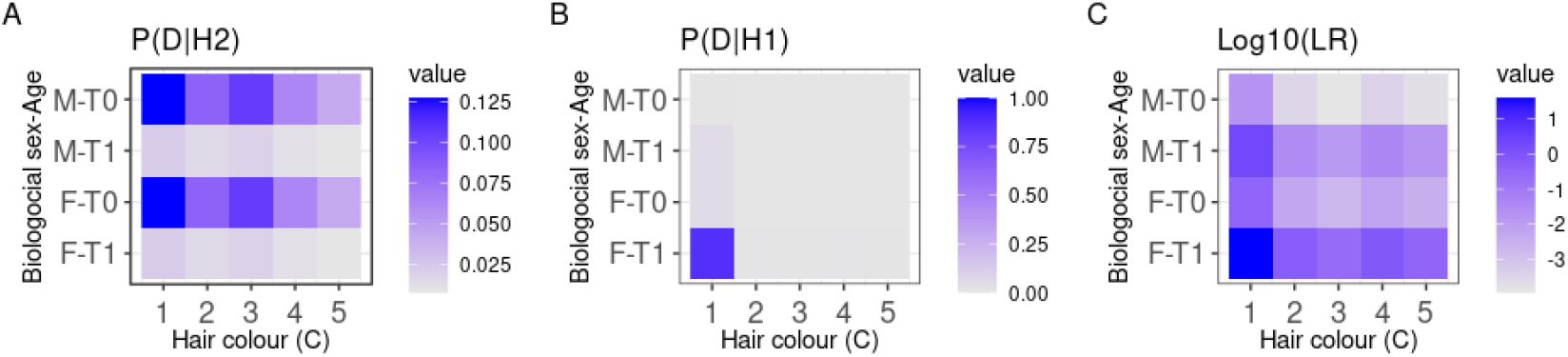
General description: This figure represents the phenotype probability distributions conditioned by the reference population (A) and the specific MPs (B). Also, the LR distribution (C) is presented. The variables are the combination of Sex and Age in the vertical axis (these are independent and are shown together in the matrix just for easy visualization): F-T0, female (*S*_*UP*_ = *f*) with an Age mismatch (*M* = 0), F-T1, female (*S*_*UP*_ = *f*) with an Age match (*M* = 1), M-T0 male (*S*_*UP*_ = *m*) with an Age mismatch (*M* = 0) and M-T1 male (*S*_*UP*_ = *m*) with an Age match (*M* = 1). Note that C could be obtained doing Log10(B/A).

**Figure 5.**
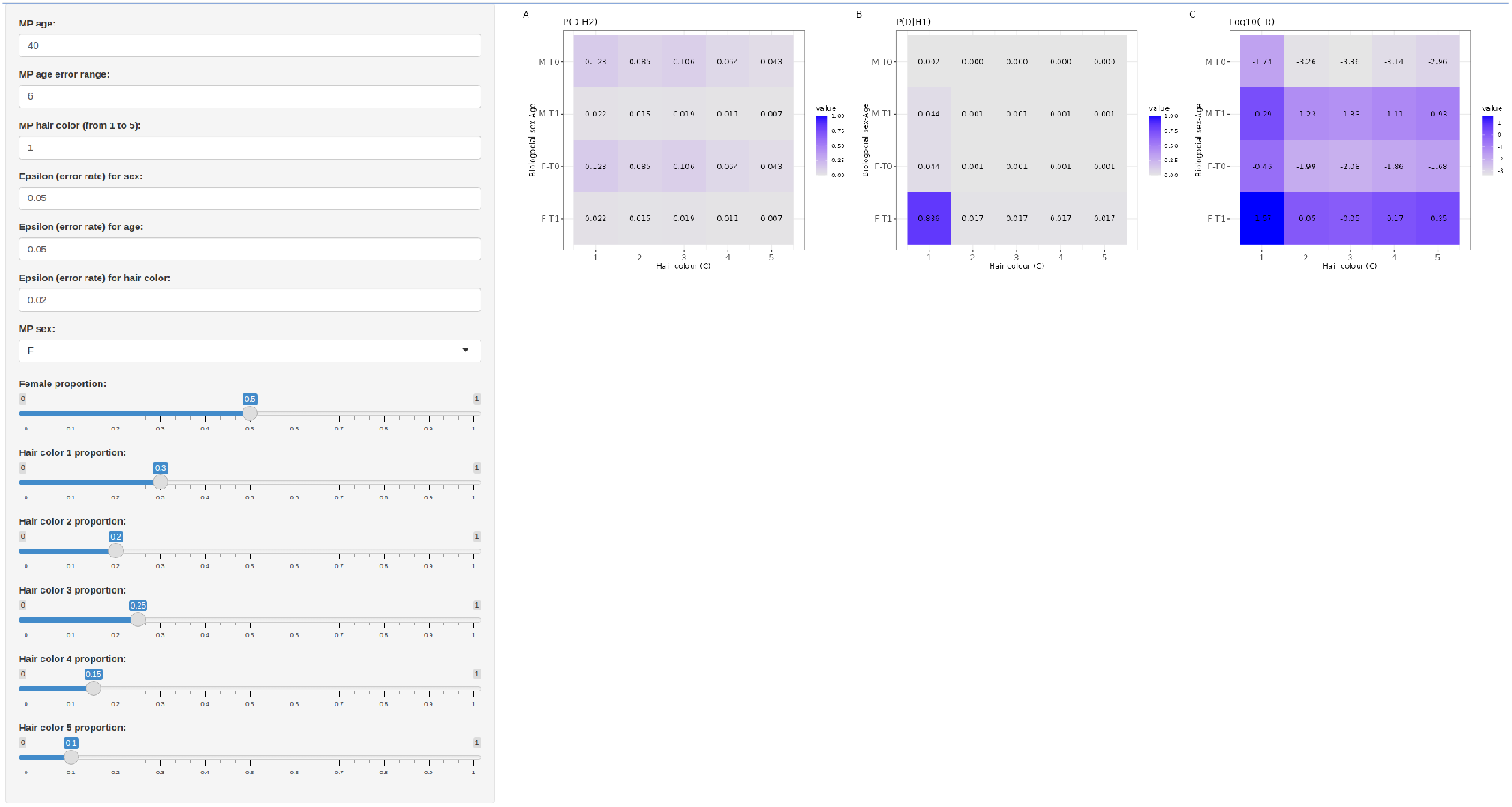
Screenshot of mispiApp, showing the sidebar with the options for the different parameters and the likelihoods and LR distribution tables. Iteratively, the user can change parameters and analyze how the likelihoods and LR change.

~~~
klPIE(POPl, MPl)
In this case, we obtain:
~~~

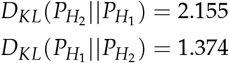

The direct interpretation of the *KL* divergence in this context is as follows: for 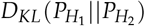, the *Log*_10_(*LR*) is expected when *H*_1_ is true; and for 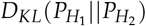, the *Log*_10_(*LR*^−1^) is expected when *H*_2_ is true. Therefore, having higher *KL* values in both directions implies an increasing difference between the mean *Log*_10_(*LR*) values expected, lower for *H*_2_ and higher for *H*_1_.

In this sense, forensic professionals might be tempted to explore different values for both parameters and characteristics and study how they affect performance. For this reason, **mispitools** incorporates a user-friendly interface that allows them to interactively change the values and analyze the tables present in Figure 4. To launch the interface, the following command must be executed:

~~~
mispiApp()
~~~

Moreover, as previously mentioned, conditional probability tables can be used to perform simulations of expected LR distributions considering *H*_1_ and *H*_2_ as true. Assuming mutual independence between DNA STR markers, sex, age, and hair color, the LRs can be combined given that the same hypotheses are tested. LR simulations can be performed with the following command:

~~~
sex_H1 <-LRsex(“F”, H = 1, LR = TRUE, seed = 123, nsims = 500)
sex_H2 <-LRsex(“F”, H = 2, LR = TRUE, seed = 123, nsims = 500)
col_H1 <-LRcol(1, H = 1, LR = TRUE, seed = 123, nsims = 500)
col_H2 <-LRcol(1, H = 2, LR = TRUE, seed = 123, nsims = 500)
age_H1 <-LRage(40, H = 1, LR = TRUE, seed = 123, nsims = 500)
age_H2 <-LRage(40, H = 2, LR = TRUE, seed = 123, nsims = 500)
~~~

Once simulated, we can combine the LRs obtained from the PIE with those calculated in the previous example (only FDE). In order to simplify the example, we will assume that MP in Ped1 and MP in Ped2 share the same characteristics, and the reference population is the same, therefore the same non-genetic LR distribution can be used with both examples. That is, we can compute the combined LRs as follows:

~~~
datasimx2 <-simLR2dataframe(datasimx)
combinedx_H1 <-datasimx2$Related * sex_H1$LRs * col_H1$LRc * age_H1$LRa
combinedx_H2 <-datasimx2$Unrelated * sex_H2$LRs * col_H2$LRc * age_H2$LRa
combined_datasimx <-as.data.frame(cbind(combinedx_H2, combinedx_H1))
names(combined_datasimx) <-c(“Unrelated”, “Related”)
combined_datasimx
~~~

~~~
Unrelated        Related
8.109142e-04 142668.8035
5.033040e-07   3739.3775
1.322667e-11   2774.3613
3.567114e-06  92513.6636
5.578832e-06    361.9391
1.631195e+00  22997.6685
~~~

For Ped2, we will proceed in a similar way:

~~~
datasimy2 <-simLR2dataframe(datasimy)
combinedy_H1 <-datasimy2$Related * sex_H1$LRs * col_H1$LRc * age_H1$LRa
combinedy_H2 <-datasimy2$Unrelated * sex_H2$LRs * col_H2$LRc * age_H2$LRa
combined_datasimy <-as.data.frame(cbind(combinedy_H2, combinedy_H1))
names(combined_datasimy) <-c(“Unrelated”, “Related”)
combined_datasimy
~~~

~~~
Unrelated       Related
1.188810e-02 672.604173
1.171118e-03   6.638100
9.179705e-05   8.274109
7.344176e-03  35.083415
4.571806e-03  88.665734
8.449938e-02   4.778942
~~~

After combining the LRs, we can use the combined_datasimx and combined_datasimy as we did in the previous example with datasimx and datasimy. This means that we can plot the LR distributions, error rates per threshold, and DT. For comparison purposes, we will show the results of the performance analyzes and DT selection.

~~~
DeT(combined_datasimx, 10)
“Decision threshold is: 3”
Trates(combined_datasimx, 3)
“FNR = 0.04 ; FPR = 0 ; MCC = 0.960768922830523”
DeT(combined_datasimy, 10)
“Decision threshold is: 6”
Trates(combined_datasimy, 6)
[1] “FNR = 0.048 ; FPR = 0 ; MCC = 0.953098602750463”
~~~

It can be demonstrated how adding LRs based on non-genetic information results in a performance improvement. Finally, if we have multiple pigmentation traits, we can use the model that takes into account the conditional dependency between them. This implies having parameters and population data for all three variables: hair, eye, and skin color. This can be evaluated with the following function:

~~~
data <-simRef()
conditioned <-conditionedProp(data, 1, 1, 1, 0.01, 0.01, 0.01)
unconditioned <-refProp(data)
likelihoods <-compute_LRs_colors(conditioned, unconditioned)
LRcols <-LRcolors(likelihoods)
Unrelated       Related
8.345434e-04 40.4032073
4.040321e+ 01 1.7348387
4.165279e-05 0. 8674194
4.040321e+01 40.4032073
1.387871e+00 40.4032073
4.081974e-01  0.1309312
~~~

These values can substitute the LR based on a unique color, such as the following:

~~~
datasimx2 <-simLR2dataframe(datasimx)
combinedx_H1 <-datasimx2$Related * sex_H1$LRs * LRcols$Related * age_H1$LRa
combinedx_H2 <-datasimx2$Unrelated * sex_H2$LRs * LRcols$Unrelated * age_H2$LRa
combined_datasimx2 <-as.data.frame(cbind(combinedx_H2, combinedx_H1))
names(combined_datasimx2) <-c(”Unrelated”, ”Related”)
combined_datasimx2
Unrelated         Related
2.087076e-07 1777703.1042
3.135672e-04    2000.6577
2.831768e-14     742.1753
2.222374e-03 1152752.5498
2.653165e-04    4509.8880
2.053480e-01     928.6272
~~~

Given these values, we can compute the DT and performance, now considering both PIE and FDE.

~~~
DeT(combined_datasimx2, 10)
“Decision threshold is: 3”
Trates(combined_datasimx2, 3)
“FNR = 0.02 ; FPR = 0 ; MCC = 0.980196058819607”
~~~

Here we can analyze and quantify the improvement in the search performance of adding PIE based LRs.

## Summary

This paper introduces an R package of missing person search statistical tools named **mispitools**. The package is available on the Comprehensive R Archive Network (CRAN) at http://CRAN.R-project.org/package=mispitools. It contains functions to calculate likelihood ratios (LRs) based on genetic data (FDE) or preliminary investigation data (PIE). In addition, it can perform simulations of the databases for the search of missing persons. This package implements previously developed methodologies for likelihood ratio threshold, error rates, and MCC coefficient estimations. It also incorporates new functions, allowing the methodologies to be combined. The functions are described and implemented using two examples. Through these examples, we show that the incorporation of likelihood ratios based on nongenetic data can be useful in large-scale missing person searches, especially when hindered by low statistical power.

## Acknowledgements

The development of these methods has been greatly influenced by extensive discussions with colleagues and professionals in forensic investigations. We particularly acknowledge Thore Egeland and Magnus Vigeland for their insightful discussions and contributions to open-source forensic packages, which are integral to this work. We are also deeply grateful to Inés Caridi and Ariel Salgado for their invaluable input, which significantly improved the conceptual and practical aspects of the methods. Furthermore, we extend our thanks to all contributors who provided suggestions and pull requests to the open source **mispitools** project, with special recognition given to Martin Amigo, Alina Hordiienko, Yike Cheng, Ozcan Mirray and Maja Ostrowska.

