## Supplementary code for "Mispitools: An R package for comprehensive statistical methods in Kinship Inference": marsico.R

```
library(devtools)
load_all()
document()


# Setting up

install.packages("mispitools")
install.packages("pedtools")
library(mispitools)
library(pedtools)
library(forrel)


# Example 1

set.seed(1234)
f <- getfreqs(Argentina)
ped1 <- linearPed(3)
ped1 <- setMarkers(ped1,locusAttributes = f)
ped1 <- profileSim(ped1,N = 1,ids = c(2,4))
ped2 <- cousinPed(1)
ped2 <- setMarkers(ped2,locusAttributes = f)
ped2 <- profileSim(ped2,N = 1,ids = 8)
datasimx = simLRgen(ped1, missing = 7, numsims = 500, seed = 1234)
datasimy = simLRgen(ped2, missing = 7, numsims = 500, seed = 1234)

plot(ped1, hatched = typedMembers(ped1))
plot(ped2, hatched = typedMembers(ped2))


LRdist(datasimx)

deplot(datasimx)

LRdist(datasimy)

deplot(datasimy)


DeT(datasimx, weight = 10)
Trates(datasimx, threshold = 4)
DeT(datasimy, weight = 10)
Trates(datasimy, threshold = 5)

# Example 2

POPl <- CPT_POP(propS = c(0.5, 0.5),
                MPa = 40,
                MPr = 6,
                propC = c(0.3, 0.2, 0.25, 0.15, 0.1))
POPl


MPl <- CPT_MP(MPs = "F", MPc = 1,
              eps = 0.05, epa = 0.05,
              epc = Cmodel())
MPl


MPl/POPl

CondPlot(MPl,POPl)

mispiApp()

sex_H1 <- LRsex("F", H = 1, LR = TRUE, seed = 123, nsims = 500)
sex_H2 <- LRsex("F", H = 2, LR = TRUE, seed = 123, nsims = 500)
col_H1 <- LRcol(1, H = 1, LR = TRUE, seed = 123, nsims = 500)
col_H2 <- LRcol(1, H = 2, LR = TRUE, seed = 123, nsims = 500)
age_H1 <- LRage(40, H = 1, LR = TRUE, seed = 123, nsims = 500)
age_H2 <- LRage(40, H = 2, LR = TRUE, seed = 123, nsims = 500)

datasimx2 <- simLR2dataframe(datasimx)
combinedx_H1 <- datasimx2$Related * sex_H1$LRs * col_H1$LRc * age_H1$LRa
combinedx_H2 <- datasimx2$Unrelated * sex_H2$LRs * col_H2$LRc * age_H2$LRa
combined_datasimx <- as.data.frame(cbind(combinedx_H2, combinedx_H1))
names(combined_datasimx) <- c("Unrelated", "Related")
head(combined_datasimx)

datasimy2 <- simLR2dataframe(datasimy)
combinedy_H1 <- datasimy2$Related * sex_H1$LRs * col_H1$LRc * age_H1$LRa
combinedy_H2 <- datasimy2$Unrelated * sex_H2$LRs * col_H2$LRc * age_H2$LRa
combined_datasimy <- as.data.frame(cbind(combinedy_H2, combinedy_H1))
names(combined_datasimy) <- c("Unrelated", "Related")
head(combined_datasimy)


DeT(combined_datasimx, 10)
Trates(combined_datasimx, 3)

DeT(combined_datasimy, 10)
Trates(combined_datasimx, 6)


data <- simRef()
conditioned <- conditionedProp(data, 1, 1, 1, 0.01, 0.01, 0.01)
unconditioned <- refProp(data)
likelihoods <- compute_LRs_colors(conditioned, unconditioned)
LRcols <- LRcolors(likelihoods)
head(LRcols)


datasimx2 <- simLR2dataframe(datasimx)
combinedx_H1 <- datasimx2$Related * sex_H1$LRs * LRcols$Related * age_H1$LRa
combinedx_H2 <- datasimx2$Unrelated * sex_H2$LRs * LRcols$Unrelated * age_H2$LRa
combined_datasimx2 <- as.data.frame(cbind(combinedx_H2, combinedx_H1))
names(combined_datasimx2) <- c("Unrelated", "Related")
head(combined_datasimx2)


DeT(combined_datasimx2, 10)
Trates(combined_datasimx2, 3)
```
